# Bacterial communities associated with the surfaces of the fresh fruits sold around Dhaka Medical College and Hospital and their anti-microbial profiles

**DOI:** 10.1101/2022.08.26.505509

**Authors:** Rubaiya Binte Kabir, Rizwana Zaman, Noor – E– Jannat Tania, Md. Asaduzzaman, Azmeri Haque, Farjana Binte Habib, Nusrat Noor Tanni, Maherun Nesa, Akteruzzaman Chowdhury, Md. Faizur Rahman, Avizit Sarker, Kakali Halder, Nazmun Sharmin, Mahbuba Chowdhury, Sultana Shazeda Nahar, Md. Mizanur Rahman, Sazzad Bin Shahid, SM Shamsuzzaman

**Affiliations:** Department of Microbiology, Dhaka Medical College, Dhaka, Bangladesh

**Keywords:** antibiogram, bacteria, Bangladesh, fresh fruits, fruit venders around hospital

## Abstract

**Background:** Now-a-days, fresh fruits are popular sources of healthy diets with low energy density. Since they are consumed raw, it may act as a source of foodborne disease and a reservoir for antibiotic resistant organisms. This study aimed to determine microbial prevalence among the fruits sold outside Dhaka Medical College Hospital (DMCH) along with their antimicrobial profiles and to detect antimicrobial resistance genes among the resistant organisms.

**Methods:** Thirty-five different types of fruits were bought from around DMCH and analyzed for the presence of bacteria. Antibiotic sensitivity was done and ESBL, AmpC β-lactamase, and MBL positive strains were identified by standard methods followed by PCR to detect ESBL, AmpC β-lactamase, and MBL genes.

**Findings:** Twenty seven different organisms were isolated and identified which were-*Klebsiella spp* (33·33%), *Citrobacter spp* (29·64%), *Enterobacter spp* (22·22%), *Escherichia coli* (11·11%), and *Staphylococcus aureus* (3·70%). Among them 48·15% organisms were resistant to different antibiotics. Only one organism (*Citrobacter spp*) produced ESBL phenotypically (7·69%). Two (15·38%) were positive for AmpC β-lactamase and one of these (*Enterobacter spp*) possessed both SHV and CTX-M15A genes by PCR. Imipenem resistance was 84·62% of the antibiotic resistant organisms, and 10 (90·91%) were phenotypically MBL positive by CD test, DDS test, and MHT. By PCR, one *Enterobacter spp* had MBL encoding gene OXA-48.

**Interpretation:** Fresh fruits, contaminated with pathogens, might be a source of transmission of resistant organisms and attribute to public health issues.

**Funding:** The office of Directorate General of Health Service, Ministry of Health and Family Welfare, Bangladesh.

**Importance:** Fruits sold around Dhaka Medical College and Hospital (DMCH) are usually consumed by patients, patient’s attendant and Health care worker. This investigation emphasized not only on the extent of bacterial contamination but also attempted to measure the anti-microbial profile of these organisms isolated from the fruits commonly available and readily consumed in and around DMCH.

## Introduction

Fresh fruits are good sources of vitamins, minerals, phytonutrients and dietary fiber. It is also considered as a measure to decrease heart diseases and some cancers.^1^ Moreover, fruits are regarded microbiologically safer than other unprocessed foods. Thus, to promote fruits and fresh products intake, the Food and Agriculture Organization (FAO) of the United Nations and the World Health Organization (WHO) recommended a minimum consumption of 400 g of vegetables and fruits per day.^2^

But, fruit surfaces are not free from microorganisms. Rather, they may carry abundance of microorganisms which may introduce new linages of pathogenic bacteria into human gut when consumed raw.^3^ These organisms mostly originate from enteric environments.^4^ As a result, fresh fruits can act as a mean of human exposure to antibiotic-resistant bacteria as well as a reservoir of antibiotic resistance genes.^5^ It certainly poses a potential public health threat since they are able to exchange resistance genes with intestinal bacteria during their colonization and passage through the intestines leading to further dissemination in the environment.^6^

Akhter *et al*. demonstrates high bacterial diversity on the fruit surfaces among which the gram negative bacteria like *Escherichia coli, Klebsiella pneumoniae, Serratia marcescens, and Pseudomonas aeruginosa and* gram positive bacteria *Bacillus cereus, Staphylococcus aureus* etc. are dominating.^7^ Extended-spectrum β-lactamase, cephalosporinase, carbapenemase and mcr gene-producing gram-negative bacteria isolated from fresh vegetables and fruit have been reported in several studies.^8,9^

The transfer of these organisms to fresh fruits and vegetables may occur at multistage from production to home kitchen-during production through use of animal manure and contaminated irrigation water, during the post-harvest stage, transport, conservation, and processing by handlers.^10^

Therefore, in this study, we determined the extent of bacterial contamination and an anti-microbial profile of the organisms isolated from the fruit surfaces, sold, and consumed in and around DMCH.

### Materials & Methods

This cross sectional study was conducted at the department of Microbiology, Dhaka Medical College, Dhaka from April 2021 to September 2021 in 5 phases at different time. Thirty five different raw fruits, namely, five apples, five mangoes, five guava, five sets of blackberry, five bunches of grapes, five oranges, four pineapples, and one sugarcane juice, were bought from the fruit venders around Dhaka Medical College and hospital, Dhaka. Samples were transported to the laboratory carefully in sterile polythene bags for bacteriological analysis.^11^ Fruits were washed with 20 ml sterile distilled water and fluid was transferred into separate beaker. A ten-fold serial dilution of the samples was performed in the Trypticase Soya broth.

### Determination of total viable count (TVC) by bacterial enumeration

Spread plate method was used for bacterial enumeration to determine the number of colony forming units (CFUs).^12^

### Isolation and identification of bacteria

After incubating the diluted samples in TSB for 24 hours, they were subcultured in blood agar and MacConkey agar media followed by incubation at 37°C for 48 hours. A single colony was further subcultured until pure culture was obtained. Identification of bacteria was performed on the basis of colony morphology, Gram’s staining reaction, and biochemical tests (catalase, coagulase, sugar utilization, gas and H2S production in Triple Sugar Iron agar media, citrate utilization test in Simmon’s Citrate agar, motility, indole and urease production in Motility Indole Urea agar media).^13^

### Antimicrobial susceptibility test

Susceptibility to antimicrobial agents of all isolated organisms were determined by modified Kirby-Bauer technique using Mueller-Hinton agar media. Zones of inhibition were interpreted according to CLSI guidelines.^14^ FDA guideline were followed for tigecycline.^15^

### Detection of ESBL producers by double disc synergy test

30µg ceftazidime disc and amoxiclav (20µg + 10µg) disc were placed 20-25mm apart (center to center) in the Mueller-Hinton agar plate and was observed for clear extension of the edge of zone of inhibition of cephalosporin disc towards amoxyclav disc after incubating at 37°C for 24 hours.^16^

### Phenotypic detection of carbapenemase producers

Organisms were considered carbapenemase producer if at least positive for two or more methods.

### Double disc synergy (DDS) test

After inoculating the test inoculums (compared with McFarland standard) in Mueller-Hinton agar plates, imipenem disc was placed on the inoculated plate along with a blank disc containing 20µl of Tris-EDTA and 20µl of 1: 320 diluted 2-mercaptopropionic acid, placed 10mm apart. A clear extension of the edge of the inhibition zone of imipenem disc towards Tris-EDTA-MPA disc was observed after incubating at 37°C for 24 hours.^17^

### Combined disc (CD) assay

Two imipenem discs (one supplemented with 5µl of 0·5 M EDTA solution containing approximately 930µg EDTA) were placed on an inoculated Mueller-Hinton agar plate following incubation at 37°C for 24 hours. An increased zone of diameter of ≥6mm around the disc containing imipenem supplemented with EDTA compared to the disc containing imipenem only was interpreted as MBL producer.^18^

### Modified Hodge test (MHT)

A lawn culture of 1:10 dilutions of 0·5 McFarland’s standard *Escherichia coli* ATCC 25922 broth was done on a Mueller-Hinton agar plate. A 10µg imipenem disc was placed in the center of the plate. Then 0·5 McFarland’s standard were made by three imipenem resistant gram negative organisms and streaked from the edge of the disc to the periphery of the plate in three different directions. After overnight incubation, the presence of clover leaf shaped zone of inhibition was interpreted as MHT positive.^19^

### Detection of anti-microbial resistance genes by PCR

#### DNA extraction

300μl distilled water was mixed with bacterial pellets and vortexed until mixed well. Then DNA was extracted in block heater (DAIHA Scientific, Seoul, Korea) at 100°C for 10 minutes for boiling followed by cooling on ice pack and centrifugation at 4°C at 13500g for 10 minutes. The extracted DNAs were kept at -20°C for future use.

#### Amplification through thermal cycler

PCR was performed in a DNA thermal cycler (Eppendorf AG, Mastercycler gradient, Hamburg, Germany) after mixing of mastermix and primers with DNA template. Each PCR run comprised of preheat at 94°C for 10 minutes followed by 36 cycles of denaturation at 94°C for 1 minutes, annealing at specified temperatures for 45 seconds, extension at 72°C for 1 minutes with final extension at 72°C for 10 minutes. Gel electrophoresis was done in 1·5% agarose (Bethesda Research Laboratories). DNA bands were detected by staining with ethidium bromide (0·5 μl/ ml) for 30 minutes at room temperature and visualized with UV transilluninator (Gel Doc, Major Science, Taiwan).

#### Data analysis

The result of the study was recorded systematically. Data analysis was done by using ‘Microsoft Office Excel 2010’ program and according to the objectives of the study. The test of significant was calculated by using X^2^ test. P value < 0·05 was taken as minimal level of significance.

## Results

Fruits were collected on five occasions and 35 cultures were done. Among those 35 fruits 25 (71·43%) yielded growth of 27 different gram positive and gram negative organisms, two fruits showed growth of double organisms. Ranges of microbial count in apple was 1×10^2^ to 6×10^2^ CFU/ml, guava was 3×10^2^ to 3×10^3^ CFU/ml, blackberry was 0·1×10^2^ to 3×10^2^ CFU/ml, orange was 0·3×10^2^ to 2·4×10^2^ CFU/ml, mango was 1·2×10^2^ to 8×10^2^ CFU/ml, grape was 1×10^2^ to 1·4×10^3^ CFU/ml and pineapple was 1·6×10^2^ CFU/ml, and sugar cane juice was 2×10^2^ CFU/ml.

Among 27 different organisms nine (33·33%) were *Klebsiella spp* eight (29·64%) were *Citrobacter spp*, six (22·22%) were *Enterobacter spp E. coli* were three (11·11%) and lastly *Staphylococcus aureus* were one (3.70%) (Table 1).

**Table 1:** Distribution of organisms associated with different fruit surfaces (N=27):

|  | Apple | Mango | Guava | Blackberry | Grapes | Orange | Pineapple | Sugar<br>cane<br>juice | Total |
| --- | --- | --- | --- | --- | --- | --- | --- | --- | --- |
|  | (N=4) | (N=5) | (N=5) | (N=5) | (N=2) | (N=4) | (N=1) | (N=1) | (N=27) |
|  |  |  |  |  |  |  |  |  | n (%) |
| *Klebsiella spp.* | 1 | 1+1* | 2+1* | .. | .. | 1 | 1 | 1 | 9 (33·33) |
| *Escherichia coli* | 1 | .. | .. | .. | 1 | 1 | .. | .. | 3 (11·11) |
| *Enterobacter*<br>*spp.* | 1 | 2 | 1 | 2 | .. | .. | .. | .. | 6 (22·22) |
| *Citrobacter spp* | 1 | 1* | 1* | 2 | 1 | 2 | .. | .. | 8 (29·64) |
| *Staphylococcus*<br>*aureus* | .. | .. | .. | 1 | .. | .. | .. | .. | 1 (3·70) |

\*- denotes presence of both organisms in the same sample.

**Table 2:**
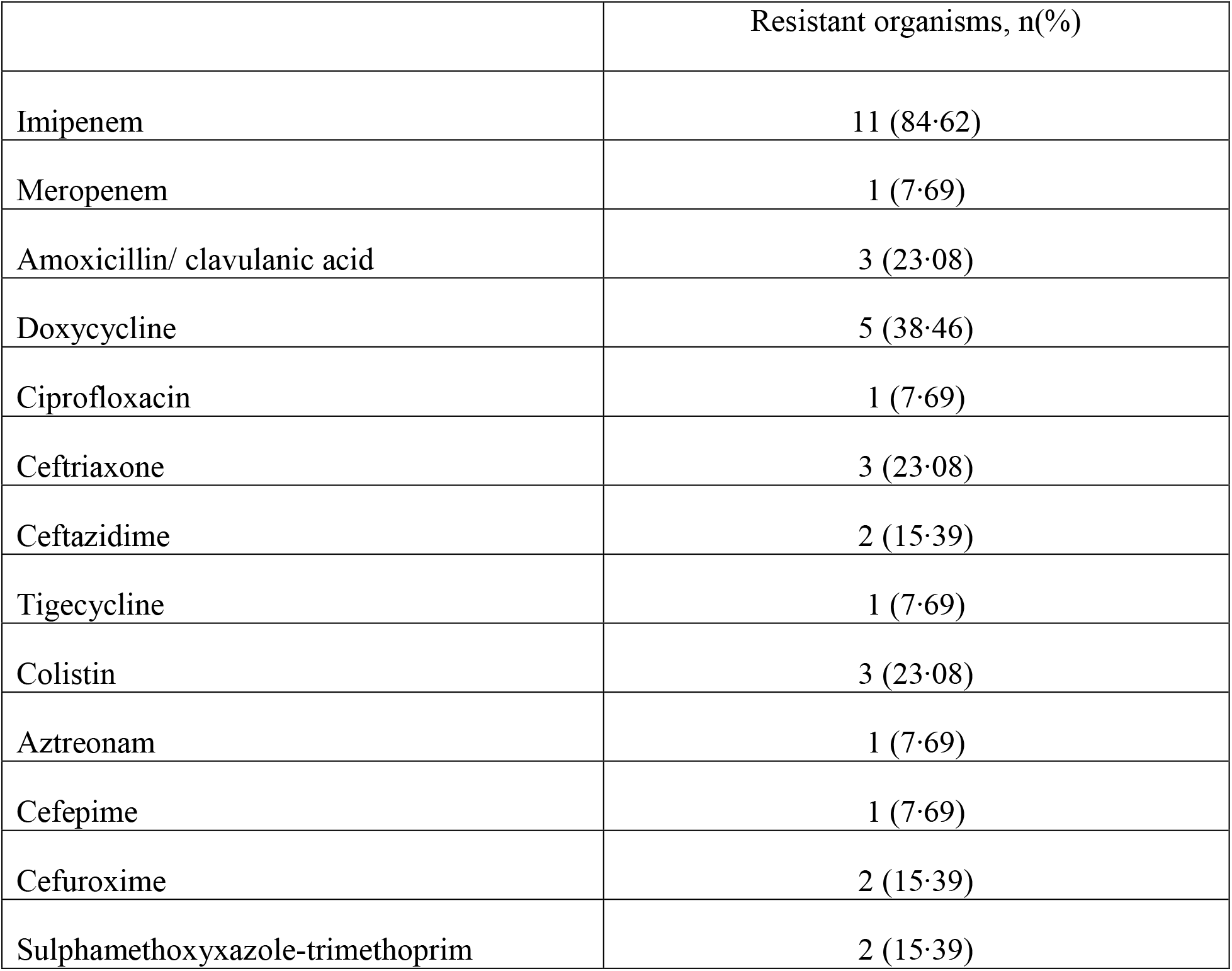
Antimicrobial resistance pattern of isolated organisms from different samples (N=13).

Fourteen (51·85%) organisms were sensitive to all drugs, and 13 (48·15%) organisms were found to have resistance to different antibiotics.

Four out of six *Enterobacter spp* and two of the nine *Klebsiella spp* were multidrug resistant. Within the resistant organisms, only one (7·69%) was extended spectrum-β-lactemase (ESBL) producer (*Citrobacter spp*) and two (15·38%) were AmpC-β-lactemase producers (*Klebsiella spp* and *Enterobacter spp*) phenotypically. One of the AmpC-β-lactemase producers (*Enterobacter spp*) was positive for both SHV and CTX-M15A gene.

Eleven out of 13 antibiotic resistant organisms (84·62%) were imipenem resistant and all of them were phenotypically detected as carbapenemase producers by CD test, DDS test or MHT. Ten (90·91%) of them were phenotypically positive for MBLs (Table 3), considering one that was only positive for MHT. Among the ten MBL positive samples, only one (*Enterobacter spp*) was positive for OXA-48 gene.

**Table 3:**
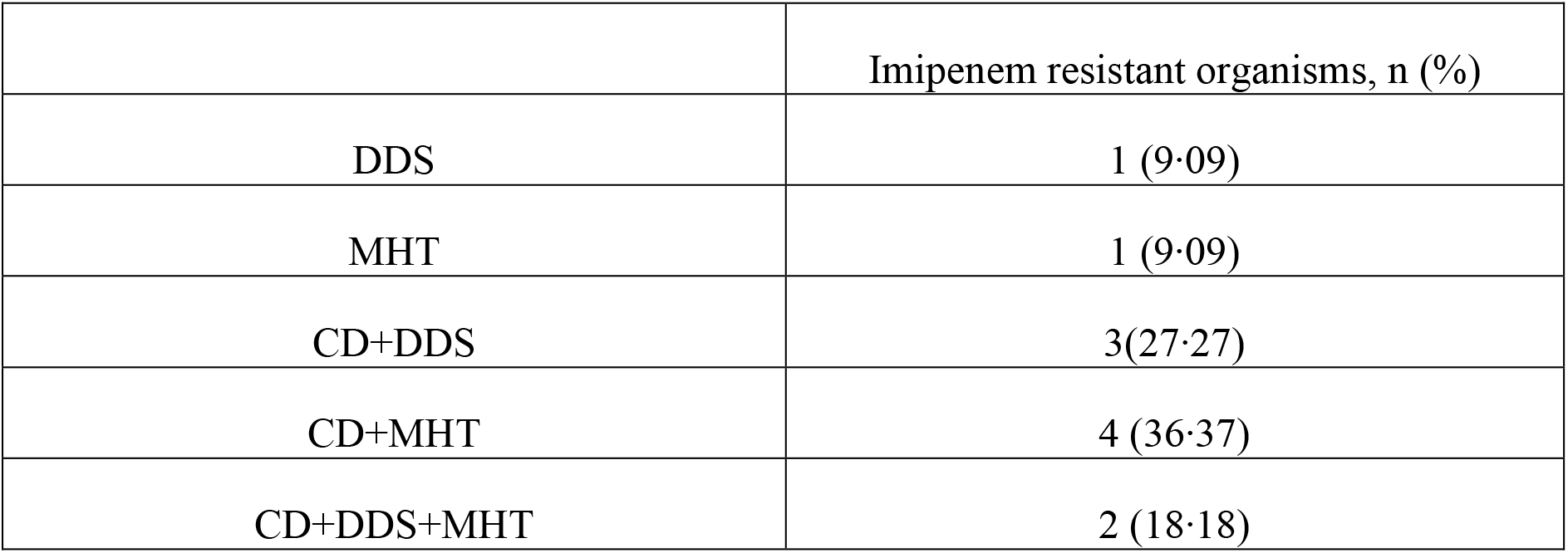
Phenotypic detection of carbapenemase producers among imipenem resistant isolates by DDS, CD and MHT (N=11).

## Discussion

Fruits are consumed in raw state which facilitates the diminished quality and health hygiene due to association of microorganisms to accumulate in a number of ways on fruit surfaces.^20^ Bacterial enumeration identified in this study highlighted the fact that fresh fruits contaminated with pathogenic organisms can act as a transmission vehicle for human diseases. It can cause a lot of sufferings for the patients, like a prolonged hospital stay, which ultimately inflects the treatment cost of the patients.

Fruits can be contaminated easily during storage and on transport.^21^ Some studies also confirmed the effect of wastewater for irrigation playing a pivotal role in contaminating fresh farm products such as fresh vegetables and fruits.^22^ In a previous study, among 25 fruit samples, 106 bacterial isolates were identified (*Escherichia coli, Salmonella spp, Vibrio spp, Bacillus spp*, and *Staphylococcus spp*).^23^ On the contrary, in this study, we identified 27 isolates of six different species among 35 samples. The difference between the findings may be due to the different samples and geographic area. Use of antibiotic in plant agriculture is in trend for quite a long time.^24^ As a result, antibiotic-resistant organisms in agricultural food is emerging throughout the world.^25^ They spread from fruits and vegetables to human via the food chain. In this study, almost half of the isolates were resistant to one or more antimicrobial groups. Among them 84·62% were resistant to imipenem, followed by 38·46% to doxycycline. 23·08% and 15·39% were resistant to third generation cephalosporin (ceftriaxone and ceftazidime). At present colistin is considered as the antibiotic of last resort and it is used when a bacteria shows resistance to most of the available antibiotics including carbapenems. In the present study, 23·08% gram negative bacilli were resistant to colistin which is a concern. Because patients of hospital consumes fresh fruits as a source of nutrients and if they get infection from fruits it may cause harm. In a study conducted in Bangladesh Agricultural University, Mymensingh, bacterial isolates of guava such as, *Escherichia coli, Vibrio spp*., and *Staphylococcus spp* were found resistant to ampicillin and cephalexin and *Salmonella spp* were resistant to chloramphenicol, ampicillin, and cephalexin.^26^ In 2018, a study on fresh vegetables revealed 83·3% prevalence of ESBL-producing strains.^27^ Whereas, in the present study, only one of the resistant isolates (*Citrobacter spp*) was positive for ESBL phenotypically. Furthermore, in this study, one AmpC β-lactamase producer harbored both SHV and CTX-M15A gene. According to a recent study, *Escherichia coli* is a predominant reservoir of CTX-M15A gene.^29^ CTX-M15A gene was also found previously in poultry, wild birds as well as in environmental water in Bangladesh.^30,31^ These findings strengthen the fact that AmpC β-lactamase gene producing organisms can cause community acquired and hospital acquired infections. In gram negative bacteria isolated from clinical samples from patients of DMCH showed increased proportion of ESBL producers up to about 2015, after that increasing trend of MBL producers was observed. The reason behind it might be due to the fact that ESBL producers are treated by carbapenems and the bacteria have started producing more carbapenemase enzymes due to increased used of carbapenems. Moreover, imported fresh food products seem to be a possible reservoir of carbapenemase-producing gram negative organisms, especially Enterobacteriaceae.^28^ In a study conducted in Algeria reported OXA-48 producing *Klebsiella pneumoniae*.^9^ In this study, we found one *Enterobacter spp* harboring OXA-48.

Wide spread use of cephalosporins might contribute in the emergence of multidrug resistant bacteria. Moreover, application of animal manure to the agricultural field can also spread drug-resistant bacteria to plant which may contribute to the contamination of fruit surfaces during collection from plant and transportation.^32^

## Conclusion

The findings of this study clearly showed that, there was a wide array of microorganisms in different fruit samples collected from around a tertiary level hospital in Bangladesh-*Klebsiella spp* was the most prevalent one. Almost half of the organisms showed resistance to multiple antibiotics and antibiotic resistance genes were found in *Enterobacter spp*. It is a matter of great concern for the consumers. It also emphasizes on the global risk of transmission of antibiotic resistant organisms via imported fresh fruits that needs to be paid attention as a public health issue.

## Acknowledgement

The authors acknowledge the Office of the Director General of Health Services (DGHS), Bangladesh for financial support and thank all the participants who volunteered for this study. They sincerely thank Principal Professor Dr. Md. Titu Miah and vice Principal Dr. Md. Shafiqul Alam Chowdhury of Dhaka Medical College, Dhaka, Bangladesh for their active cooperation.

## Authors’ contribution

All authors gave their intellectual inputs while preparing the manuscript and agreed mutually to submit this article to this journal.

|  | Research concept | Study design | Protocol writing | Responds to reviewers' comments and Defending at IRB | Developing data collection tools | Data collection | Data analysis/interpretation of results | Writing manuscript | Revise manuscript |
| --- | --- | --- | --- | --- | --- | --- | --- | --- | --- |
| Rubaiya Binte Kabir |  |  |  | NA (No human subject or other animals were not involved). | √ | √ | √ | √ |  |
| Rizwana Zaman |  |  |  |  | √ | √ |  |  |  |
| Noor-E-Jannat Tania |  |  |  |  | √ | √ |  |  |  |
| Mohammad Asaduzzaman |  |  |  |  | √ | √ |  |  |  |
| Azmeri Haque |  |  |  |  | √ | √ |  |  |  |
| Farjana Binte Habib |  |  |  |  | √ | √ |  |  |  |
| Nusrat Noor Tanni |  |  | √ |  | √ | √ | √ | √ |  |
| Maherun Nesa |  |  |  |  | √ | √ |  |  |  |
| Akteruzzaman Chowdhury |  |  |  |  | √ | √ |  |  |  |
| Md. Faizur Rahman |  |  |  |  |  |  | √ |  |  |
| Avizit Sarker |  |  |  |  |  |  | √ |  |  |
| Kakali Halder |  |  |  |  |  |  |  |  | √ |
| Nazmun Sharmin |  |  |  |  |  |  | √ |  |  |
| Mahbuba Chowdhury |  |  |  |  |  |  |  |  | √ |
| Sultana Shazeda Nahar |  |  |  |  |  |  |  |  | √ |
| Md. Mizanur Rahman |  |  |  |  |  |  |  |  | √ |
| Sazzad Bin Shahid |  |  |  |  |  |  |  |  | √ |
| SM Shamsuzzaman | √ | √ | √ |  |  |  | √ | √ | √ |

